# Multiple introductions, regional spread and local differentiation during the first week of COVID-19 epidemic in Montevideo, Uruguay

**DOI:** 10.1101/2020.05.09.086223

**Authors:** Cecilia Salazar, Florencia Díaz-Viraqué, Marianoel Pereira-Gómez, Ignacio Ferrés, Pilar Moreno, Gonzalo Moratorio, Gregorio Iraola

**Author notes:** Corresponding author: Gregorio Iraola. Laboratorio de Genómica Microbiana, Institut Pasteur de Montevideo, Montevideo 11400, Uruguay.

## Abstract

**Background:** After its emergence in China in December 2019, the new coronavirus disease (COVID-19) caused by SARS-CoV-2, has rapidly spread infecting more than 3 million people worldwide. South America is among the last regions hit by COVID-19 pandemic. In Uruguay, first cases were detected on March 13 ^th^ 2020 presumably imported by travelers returning from Europe.

**Methods:** We performed whole-genome sequencing of 10 SARS-CoV-2 from patients diagnosed during the first week (March 16^th^ to 19^th^) of COVID-19 outbreak in Uruguay. Then, we applied genomic epidemiology using a global dataset to reconstruct the local spatio-temporal dynamics of SARS-CoV-2.

**Results:** Our phylogeographic analysis showed three independent introductions of SARS-CoV-2 from different continents. Also, we evidenced regional circulation of viral strains originally detected in Spain. Introduction of SARS-CoV-2 in Uruguay could date back as early as Feb 20^th^. Identification of specific mutations showed rapid local genetic differentiation.

**Conclusions:** We evidenced early independent introductions of SARS-CoV-2 that likely occurred before first cases were detected. Our analysis set the bases for future genomic epidemiology studies to understand the dynamics of SARS-CoV-2 in Uruguay and the Latin America and the Caribbean region.

## Introduction

In December 2019, a new Coronavirus disease (COVID-19) was detected in Wuhan, China. Its causative agent, known as SARS-CoV-2, has spread rapidly causing a global pandemic of pneumonia affecting more than 3M people with more than 200,000 deaths to date [1]. Global spread of SARS-CoV-2 was primarily driven by international movement of people, since travel-associated cases of COVID-19 were reported outside of China as early as mid-January 2020 [2]. This global health emergency has deployed international efforts to apply genomic epidemiology to track the spread of SARS-CoV-2 in real time. The recent development of targeted sequencing protocols by the ARTIC Network [3], open sharing of genomic data through the GISAID (www.gisaid.org) database and straightforward bioinformatic tools for viral phylogenomics [4], provides the opportunity to reconstruct global spatio-temporal dynamics of the COVID-19 pandemic with unprecedented comprehensiveness and resolution.

South America was one of the last regions in the world to be hit by COVID-19. Indeed, first cases were confirmed in Brazil around 2 months after its emergence in China. Then, SARS-CoV-2 rapidly emerged in neighboring countries like Uruguay, that reported first cases in Montevideo, its capital city, on March 13 ^th^. Population of Uruguay is among the smallest in South America (∼3.5M), and is mainly concentrated in Montevideo and its metropolitan area (∼56% of the total population). Also, the biggest international airport of the country is placed in this metropolitan area and concentrates more than 90% of international arrivals. These characteristics makes Montevideo a suitable place to apply genomic surveillance to uncover epidemiological patterns during the early phase of the COVID-19 local outbreak in Uruguay. We therefore aimed to characterize the spatio-temporal dynamics of SARS-CoV-2 by sequencing around 10% of cases occurred during the first week of outbreak in Montevideo, allowing us to identify transmission patterns, geographic origins and genetic variation among local strains.

## Methods

### Ethics statement

Residual de-identified nasopharyngeal samples were remitted to the Institut Pasteur Montevideo, that has been validated by the Ministry of Health of Uruguay as an approved center providing diagnostic testing for COVID-19. All samples were de-identified before receipt by the study investigators.

### Sample collection, processing and sequencing

Nasopharyngeal swabs were obtained from patients residing in Montevideo. No other location data was available to researchers to avoid patient identification. Swabs were placed in virus transport media (BD Universal Viral Transport Medium) immediately after collection. Total RNA extraction was performed directly from samples (500 μL) using TRIzol reagent (Invitrogen Life Technologies, Carlsbad, CA, USA). An in-house OneStep RT-qPCR assay based on TaqMan probes targeting the N gene and the Open Reading Frame 1b region were used to test for the presence of SARS-CoV-2 RNA. We selected a total of 10 positive samples with cycle-threshold (CT) less than 30 for downstream whole-genome sequencing.

Sequencing of SARS-CoV-2 positive samples was performed according to the primalseq [5] approach on the MinION sequencing platform (Oxford Nanopore Technologies, United Kingdom) using the V2 primer pools. Sequencing libraries were prepared based on the PCR tiling of COVID-19 virus protocol (ARTIC Network) using the Ligation Sequencing Kit (SQK-LSK109) and Native Barcoding Expansion (EXP-NBD114) (Oxford Nanopore Technologies, United Kingdom), with the following modifications: i) RNA was reverse-transcribed using Superscript II reverse transcriptase (Invitrogen Life Technologies, Carlsbad, CA, USA) and random hexamer primers following the manufacturer’s instructions, ii) Phusion Hot Start II High-Fidelity PCR Master Mix (Thermo Scientific) was used for PCR amplification and, iii) Blunt/TA ligase (New England Biolabs, USA) was used to ligate barcodes to each sample. To detect cross-contamination between samples, a no template sample was added from a patient tested negative for SARS-CoV-2 from the RNA purification step.

A total of 15 ng of pooled samples was loaded onto a MinION R9.4.1 flow cell and sequenced for 5 hours generating ∼2 million reads of average quality of 12. The RAMPART software from the ARTIC Network (https://github.com/artic-network/rampart) was used to monitor the sequencing run in real time, estimate coverage across samples and check barcoding. Subsequently, basecalling was performed Guppy software v3.2.1 using the high accuracy module. Consensus genomes were generated and variants were called using the ARTIC Network bioinformatic pipeline (https://artic.network/ncov-2019/ncov2019-bioinformatic.sop.html). Amplicons that were not sequenced or whose depth was less than 20x were not included in the consensus sequences and these positions were represented by “N” stretches. See Supplementary Table S1.

### Phylogenetic and spatio-temporal analysis

To investigate the origin, dynamics and genetic variation of SARS-CoV-2 in Uruguay, we added our 10 genomes to a dataset of 2,443 complete (>29,000 bp.), high-quality genomes available from GISAID (https://www.gisaid.org) on April 2^nd^ 2020. A list of sequences and acknowledgements to the originating and submitting labs is presented in Supplementary Table S2. We analyzed this dataset using the augur toolkit version 6.4.2 [3]. Briefly, genomes were aligned using MAFFT [6] and a phylogeny was reconstructed with IQ-TREE [7]. Estimation of ancestral divergence times and geographic locations for internal nodes of the tree, and identification of branch-specific or clade-defining mutations were obtained using augur and TreeTime [8]. Code for performing these analyses can be found at http://github.com/giraola/covid-19-uruguay and results can be visualized at https://nextsrain.org/community/giraola/covid19-uruguay.

## Results

### Multiple international introductions of SARS-CoV-2 into Uruguay

To investigate the origin of SARS-CoV-2 in Uruguay we whole-genome sequenced 10 cases diagnosed with COVID-19 in Montevideo between March 16^th^ to 19^th^. This is within the first week after the presence of the virus was reported in Uruguay in March 13^th^ by the national authorities (Fig. 1). These cases represented 10.6% of accumulated cases (10 out of 94) reported in Montevideo by March 19^th^. Our phylogenetic analyses showed that genomes sequenced from cases reported in Uruguay were placed in three distinct clusters. Genome UY-10 (hereinafter referred as cluster C1-UY) was embedded within clade A1a. Two other viral genomes, UY-7 and UY-8 (hereinafter referred as cluster C2-UY) were related each other and placed within clade B1. The remaining 7 viral genomes (hereinafter referred as cluster C3-UY) were related each other and were placed within clade B (Fig. 2). These clusters were supported by specific, non-homoplasic mutations (Table 1). Together, these results indicate three independent introductions of SARS-CoV-2 into Uruguay.

**Table 1.** Cluster-defining mutations identified in SARS-CoV-2 from Uruguay.

| Cluster | Genome feature | Mutation | Type | Amino acid change |
| --- | --- | --- | --- | --- |
| C1-UY | 3'UTR | A29731G | Transition | No |
| C2-UY | ORF1ab | A11782G | Transition | No |
| C3-UY | ORF3a | C25521T | Transition | No |

**Figure 1.**
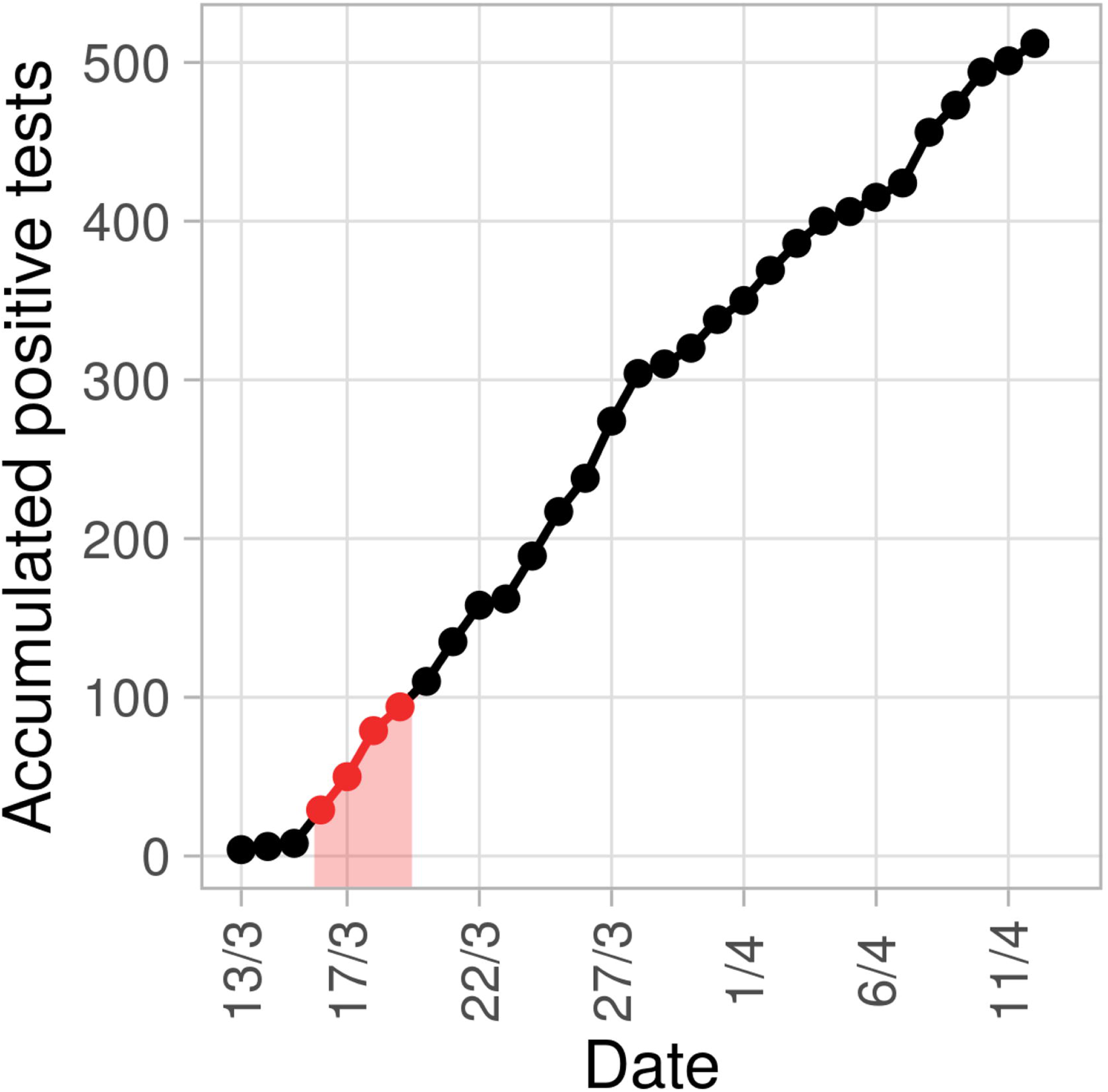
Accumulated number of COVID-19 cases in Uruguay since March 13^th^ when first cases were notified by national authorities.

**Figure 2.**
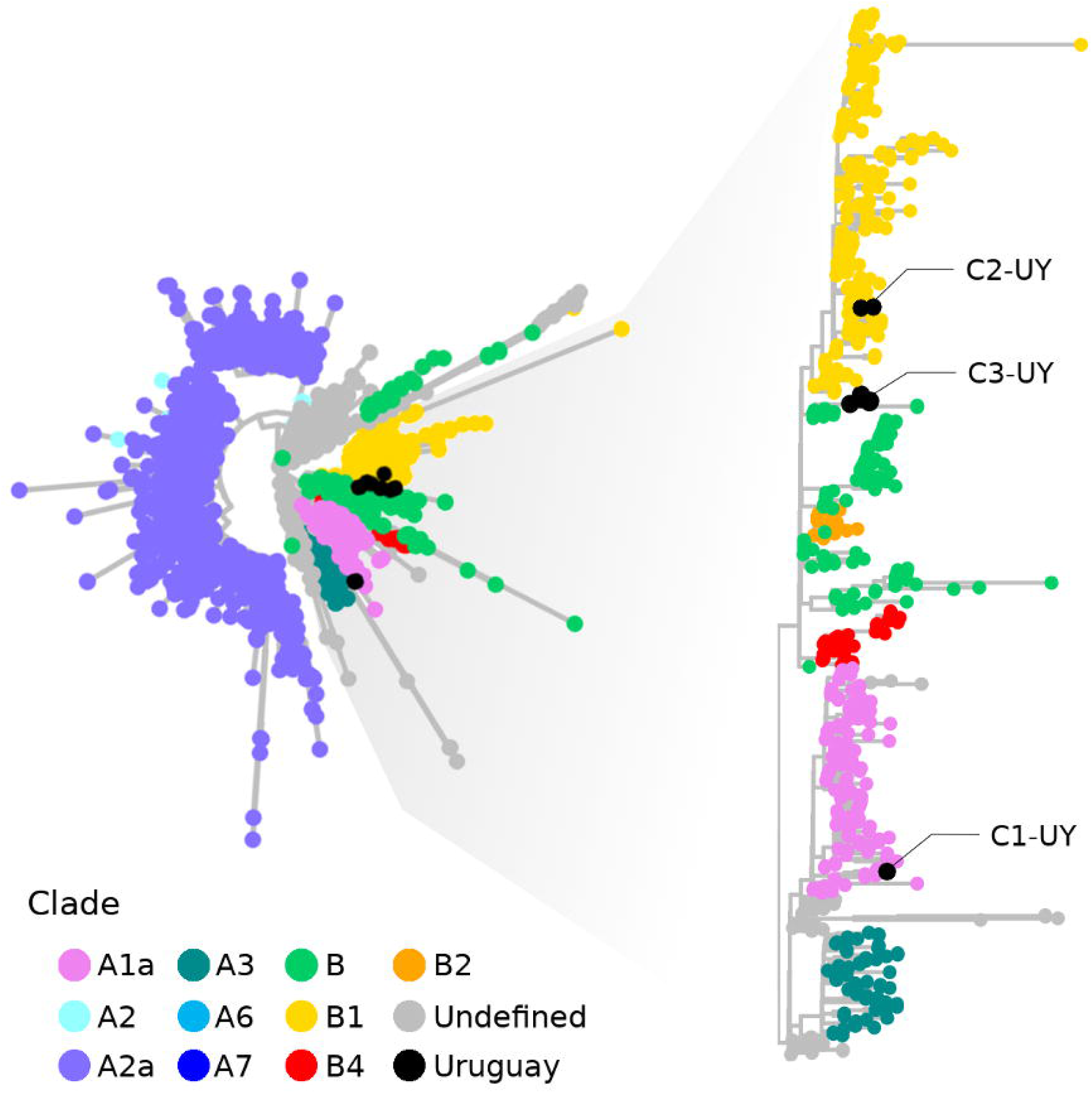
Phylogenetic relationships of 10 SARS-CoV-2 genomes from Uruguay including more than 2,500 public genomes generated worldwide and deposited in the GISAID database. Colors represent current main clades defined by Nextstrain [4].

### Early SARS-CoV-2 circulation and regional spread

To uncover the spatio-temporal dynamics of SARS-CoV-2 in Uruguay, we performed a phylogeographic reconstruction. The most probable ancestral location for C1-UY was Australia, however, very similar viruses have been also sequenced from the United Kingdom (Fig. 3). The last common ancestor of C1-UY in Australia dated back to Mar 5 ^th^ (Confidence Interval (CI): Feb 28^th^ - Mar 13^rd^). C2-UY was embedded in a cluster of viruses sequenced in Canada, the United States, Australia and Iceland whose emergence dated back to February 28^th^ (CI: Feb 25^th^ - Mar 4^th^) (Fig. 4). C3-UY likely originated from European viruses circulating in Spain since Jan 21 ^th^ (CI: Jan 7^th^ - Feb 5^th^). The last common ancestor of C3-UY in Uruguay was tracked to Mar 4^th^ (CI: Feb 23^rd^ - Mar 11^th^) (Fig. 5). Two viruses from Chile were intermingled between viruses from Spain and Uruguay, indicating regional circulation of these variants. Together, these results indicate that SARS-CoV-2 could be circulating in the country previous to its first detection in Mar 13^th^.

**Figure 3.**
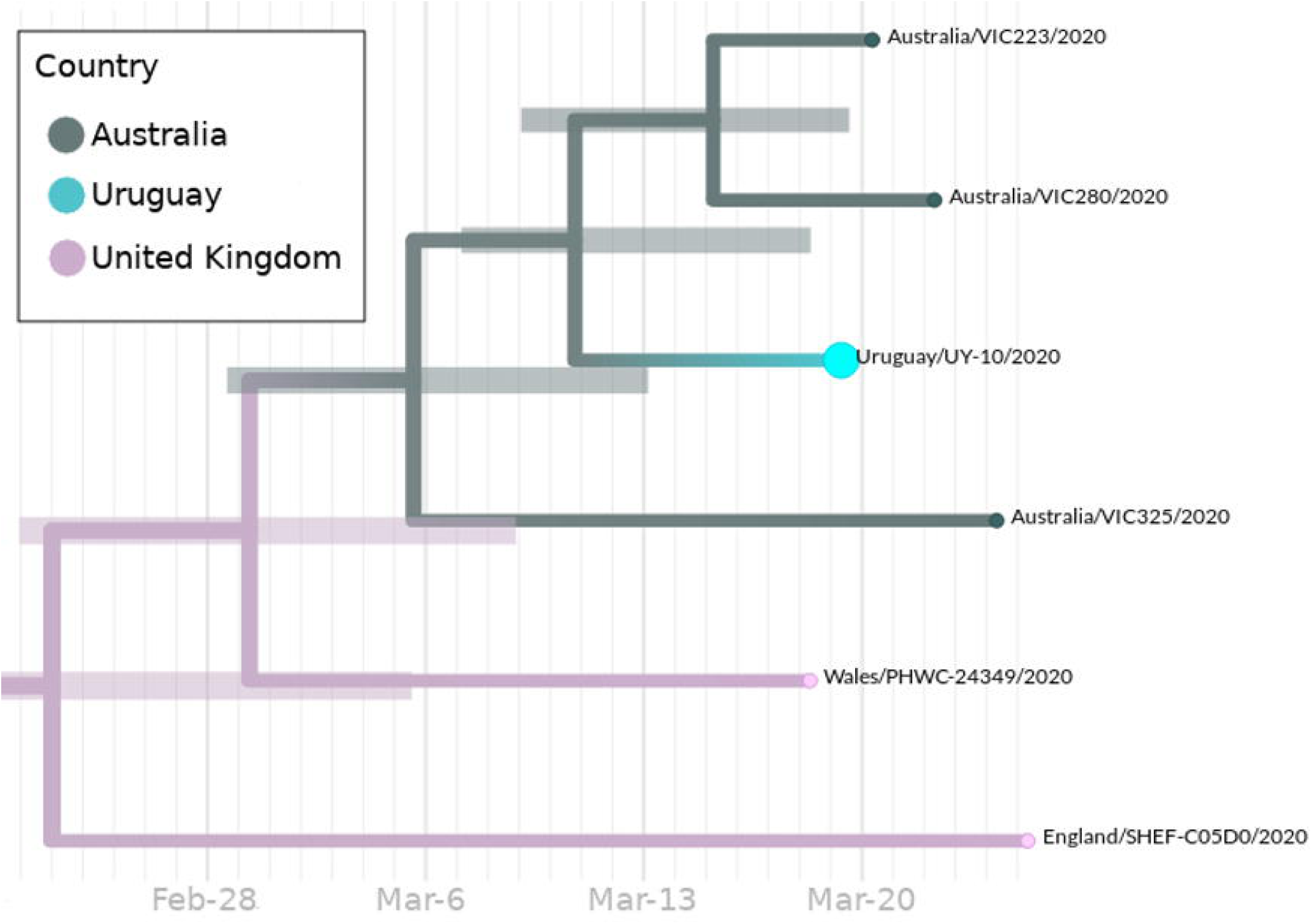
Dated phylogeny showing C1-UY. Branches and nodes are colored according to the most probable ancestral geographic location. Confidence intervals for inferred dates are shown as node bars.

**Figure 4.**
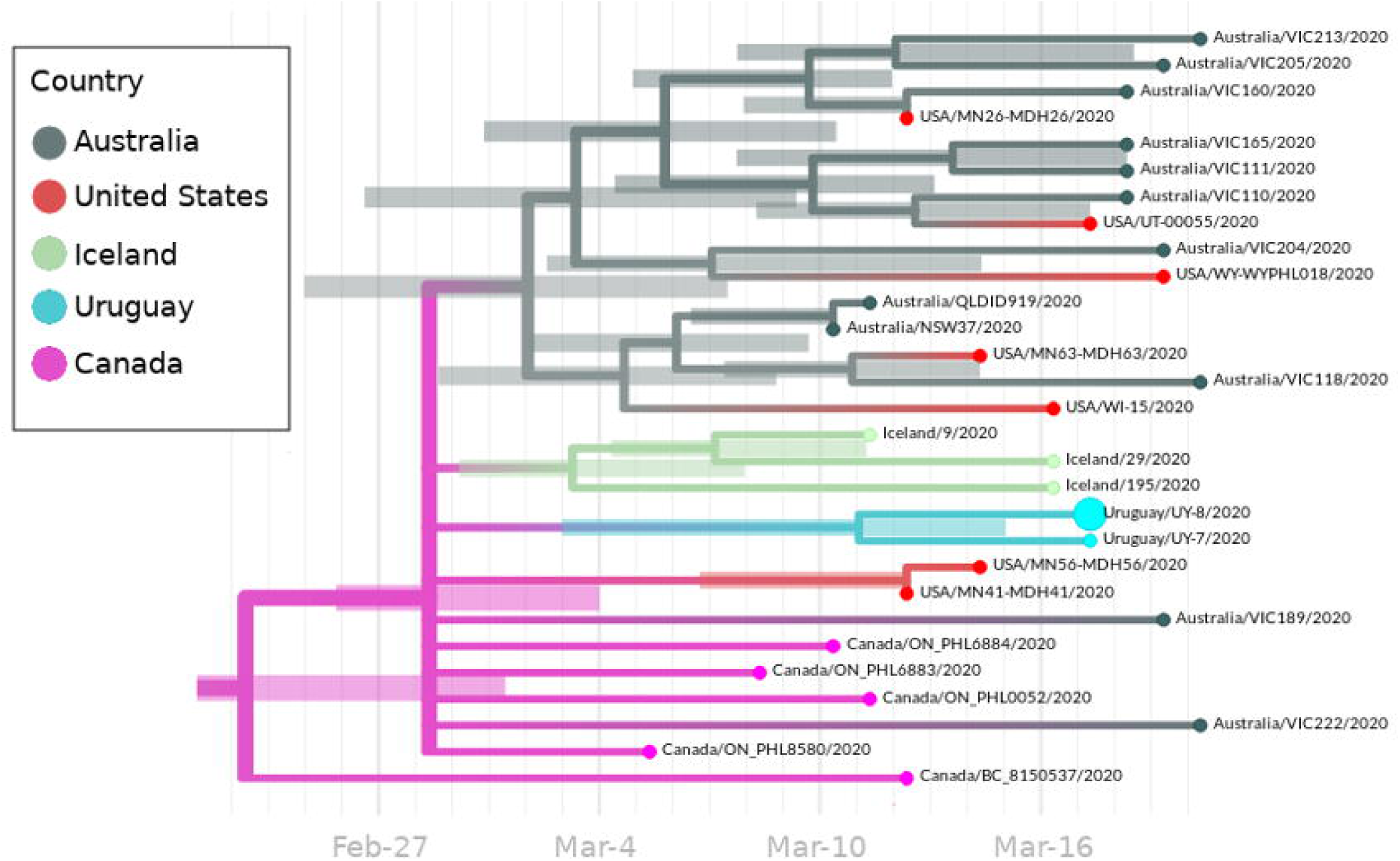
Dated phylogeny showing C2-UY. Branches and nodes are colored according to the most probable ancestral geographic location. Confidence intervals for inferred dates are shown as node bars.

**Figure 5.**
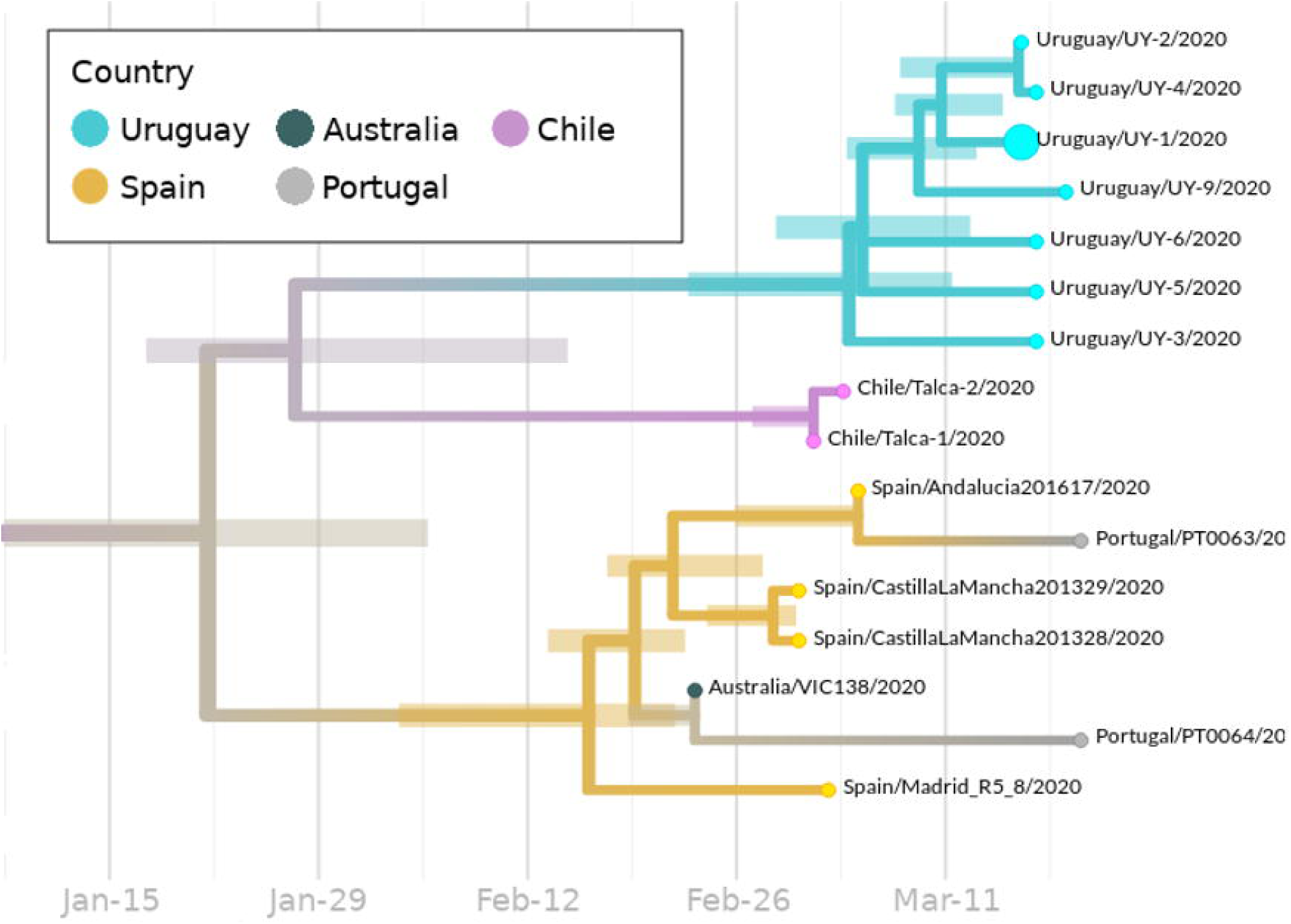
Dated phylogeny showing C3-UY. Branches and nodes are colored according to the most probable ancestral geographic location. Confidence intervals for inferred dates are shown as node bars.

### Local clusters and genetic differentiation

To further characterize Uruguayan SARS-CoV-2, we identified genetic differences among genomes. C1-UY ancestor likely located in Australia presented one non-synonymous mutation C3096T which affected the ORF1ab protein (S955L) and genome UY-10 harbored one additional synonymous mutation A29731G. C2-UY ancestor likely located in Canada was characterized by one synonymous mutation A24694T and the last common ancestor to UY-7 and UY-8 presented another synonymous change A11782G. Additionally, genome UY-8 presented one synonymous mutation C17470T. C3-UY and Chilean genomes displayed one synonymous mutations C17470T which distinguished them from the original Spanish lineage. Additionally, C3-UY differentiated from Chilean genomes by one synonymous mutation C25521T. Specifically, among C3-UY genome UY-5 presented a single synonymous mutation C6310T and genome UY-6 presented one non-synonymous mutation C7635A in the ORF1ab protein (T2457K). Overall, we observed mutations within Uruguayan clusters supporting rapid local differentiation (Table 2).

**Table 2.**
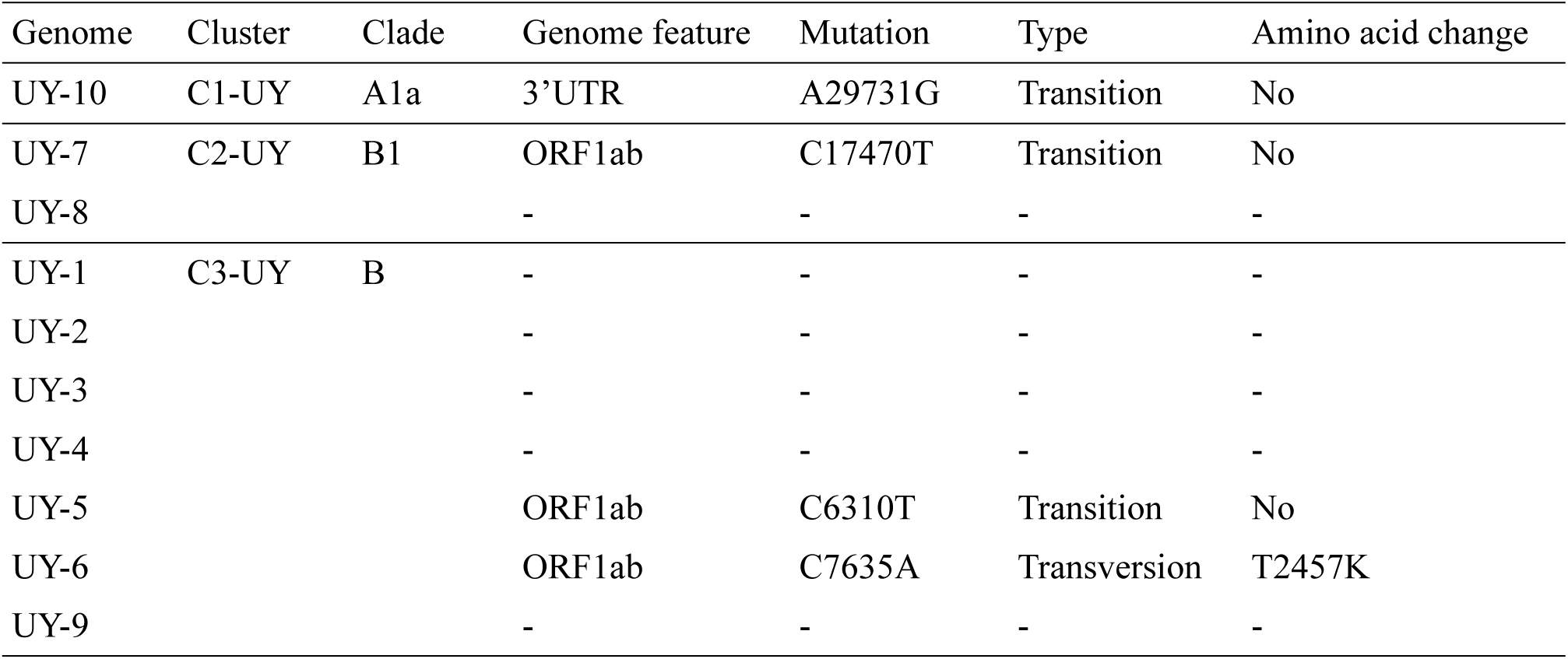
Specific mutations found in SARS-CoV-2 genomes from Uruguay.

## Discussion

Using portable, low-cost sequencing approaches based on the MinION platform (Oxford Nanopore Technologies) we were able to generate whole-genome sequences of SARS-CoV-2 just 24 hours after receiving the samples in the lab. This allowed us to determine the most plausible spatio-temporal scenario during the first week since COVID-19 was detected in Uruguay. Consequently, we show that real-time molecular epidemiology responses can be implemented locally as previously done by other countries [9].

Our results uncovered SARS-CoV-2 genomes derived from three independent international origins as early as 3 days after the disease was reported in Uruguay. International air transport has likely played a central role in the rapid spread of SARS-CoV-2, contributing to the current COVID-19 pandemic. According to air transport statistics from the World Bank (https://data.worldbank.org) and the National Government (www.dinacia.gub.uy), Uruguayan airports received ∼2.2M passengers in 2018 ranking 12^th^ within the Latin America and the Caribbean region. This highlights that even for countries with comparatively low air traffic, multiple international introductions of SARS-CoV-2 have played an important role in the establishment of local outbreaks.

We estimated these introductions could have occurred as early as mid-February, around 1 month before first cases were reported in Uruguay in March 13^th^. This is in line with observations from other geographic regions, for example, in New York first cases were confirmed in March but a recent study suggested that initial virus introductions could be traced back to February 20^th^ [10]. Also, the virus was likely introduced in Europe by a traveler that arrived from Wuhan to France, but the recent identification of a group of ill Chinese tourists in France previous to this case supports earlier introductions [11]. Indeed, SARS-CoV-2 has been proposed to circulate in humans even before its first detection in December 2019 [12].

Together, our results highlight the importance of active genomic surveillance during the ongoing COVID-19 pandemic to guide informed decisions, based on assessing the epidemiological behavior of SARS-CoV-2 in real time. The application of rapid genomic epidemiology during local outbreaks allows to identify main routes of virus introduction and estimate spatio-temporal dynamics which can be used to improve contingency measures. Remarkably, tracking the emergence of local genetic variants like those we observed in Montevideo, is important to evaluate the sensibility of molecular diagnosis and potential impacts on future anti-viral strategies. Subsequent generation of a more comprehensive genomic dataset from the ongoing outbreak in Uruguay will allow us to identify domestic transmission patterns and spatio-temporal characterization of local clusters. Additionally, setting up coordinated efforts to generate genomic data in South America will allow us to perform integrative analyses to uncover SARS-CoV-2 dynamics at the continental level.

## Acknowledgements

We thank Josh Quick and the ARTIC Network for providing SARS-CoV-2 sequencing primers and support with sequencing protocols. We also thank to everyone who openly shared their genomic data on GISAID (authors listed in Supplementary Table 2). This work is part of the

